# Chemokine receptors ACKR2 and CCR1 coordinate macrophage dynamics and mammary gland development

**DOI:** 10.1101/747360

**Authors:** Gillian J Wilson, Ayumi Fukuoka, Samantha R Love, Jiwon Kim, Marieke Pingen, Alan J Hayes, Gerard J Graham

## Abstract

Macrophages are key regulators of developmental processes, including those involved in mammary gland development. We previously demonstrated that the atypical chemokine receptor, ACKR2, contributes to control of ductal epithelial branching in the developing mammary gland by regulating macrophage dynamics. ACKR2 is a chemokine-scavenging receptor, which mediates its effects through collaboration with inflammatory chemokine receptors (iCCRs). Here we reveal that ACKR2, and the iCCR CCR1, reciprocally regulate branching morphogenesis in the mammary gland, whereby stromal ACKR2 modulates levels of the shared ligand CCL7 to control the movement of a key population of CCR1-expressing macrophages to the ductal epithelium. In addition estrogen, which is essential for ductal elongation during puberty, upregulates CCR1 expression on macrophages. The age at which girls develop breasts is decreasing, which raises the risk of diseases including breast cancer. This study presents a previously unknown mechanism controlling the rate of mammary gland development during puberty and highlights potential therapeutic targets.

**Summary:** In the mammary gland during puberty, availability of the chemokine CCL7 is controlled by a scavenging receptor ACKR2 and provides a key signal to macrophages which have the receptor CCR1. Together, this controls the timing of development.

## Introduction

Breast development (thelarche) is the first visible sign of puberty in females, and typically occurs between the ages of 8 and 13 (Merke and Cutler Jr, 1996). Globally, the age at pubertal onset is falling (de Muinck Keizer-Schrama and Mul, 2001). Early puberty is associated with an increased risk of disease in later life, including type II diabetes heart disease, and cancer (Day *et al.*, 2015). Importantly, girls who develop breasts before the age of 10 are 20% more likely to develop breast cancer (Bodicoat *et al.*, 2014). Therefore, understanding the molecular and cellular mechanisms underlying breast development is of key importance.

The mammary gland develops through branching morphogenesis, giving rise to ductal epithelial networks. In the mouse this process begins at around 3 weeks (MM Richert, KL Schwertfeger, JW Ryder, 2000), when highly proliferative structures known as terminal end buds (TEBs) form at the end of epithelial ducts and drive network formation. Supporting this process is a stromal population containing fibroblasts, extracellular matrix (ECM), adipocytes and immune cells (Wiseman and Werb, 2002). Prominent amongst the stromal immune cells are macrophages which are found throughout the gland and surrounding TEBs. Macrophages have been implicated in numerous developmental processes (Wynn, Chawla and Pollard, 2013), and mammary gland development is severely impaired in macrophage-deficient mice with altered TEB formation, ductal elongation during puberty and lobuloalveoli development in pregnancy (Pollard and Hennighausen, 1994; Gouon-Evans, Rothenberg and Pollard, 2000). Overall these studies indicate a key role for macrophages in the regulation of ductal branching in the developing mammary gland.

Macrophages are recruited in a dynamic manner into the mammary gland throughout development (Coussens and Pollard, 2011). The molecular mechanisms regulating the intra-gland movement of macrophages as they migrate to terminal end buds to mediate their developmental effects, are not currently understood and insights into these mechanisms will enhance our overall understanding of how macrophages control mammary gland development. Chemokines which comprise a family of proteins characterised by a conserved cysteine motif, are important in vivo regulators of macrophage intra-tissue dynamics. The chemokine family is subdivided into CC, CXC, XC and CX3C sub-families according to the cysteine distribution, and chemokines act through G-protein coupled receptors to mediate leukocyte migration (Nibbs and Graham, 2013). Within tissues chemokine distribution, and gradients, can be regulated by members of the atypical chemokine receptor (ACKR) family, which are 7-transmembrane spanning receptors that lack classical signalling responses to ligands and which are typically stromally-expressed (Nibbs and Graham, 2013). Therefore, together, signalling chemokine receptors and ACKRs regulate intra-tissue chemokine function and coordinate leukocyte migration.

We have a long-standing interest in one of the atypical chemokine receptors, ACKR2. ACKR2 scavenges and degrades inflammatory CC-chemokines thereby regulating their intra-tissue concentration and spatial distribution (Nibbs and Graham, 2013). Accordingly it is a key player in the resolution of the inflammatory response with implications for autoimmunity and cancer (Nibbs *et al.*, 2007; Di Liberto *et al.*, 2008; Shams *et al.*, 2017). We previously demonstrated a role for ACKR2 in regulating branching morphogenesis in the developing lymphatic system via control of macrophage dynamics around developing vessels. More recently we have shown that ACKR2 also regulates branching morphogenesis in the mammary gland and ACKR2-/- mice display precocious mammary gland development. In essence, ACKR2 deficiency results in increased levels of monocyte and macrophage attracting chemokines in the developing mammary gland and this is associated with dysregulation of macrophage numbers and accelerated branching morphogenesis. The chemokines scavenged by ACKR2 are ligands for the signalling chemokine receptors CCR1, CCR2, CCR3, CCR4 and CCR5 (Fig. 1) (Nibbs and Graham, 2013; Bachelerie *et al.*, 2014). It is likely therefore that the effects of ACKR2 on mammary gland development are indirect, and a consequence of the regulation of levels of chemokines capable of modulating macrophage function via one of these 5 receptors. Curiously, the dominant monocyte recruitment receptor, CCR2, does not control the rate of branching morphogenesis in the mammary gland (Wilson *et al.*, 2017; Jäppinen *et al.*, 2019), and mammary gland macrophages do not express CCR4 (Wilson *et al.*, 2017). Together, this suggests that the phenotype seen in ACKR2-/- mammary glands is a consequence of altered responses through CCR1, CCR3 or CCR5. The purpose of this study was to determine which of these 3 receptors is the reciprocal partner of ACKR2, in the regulation of branching morphogenesis in the developing mammary gland.

**Figure 1:**
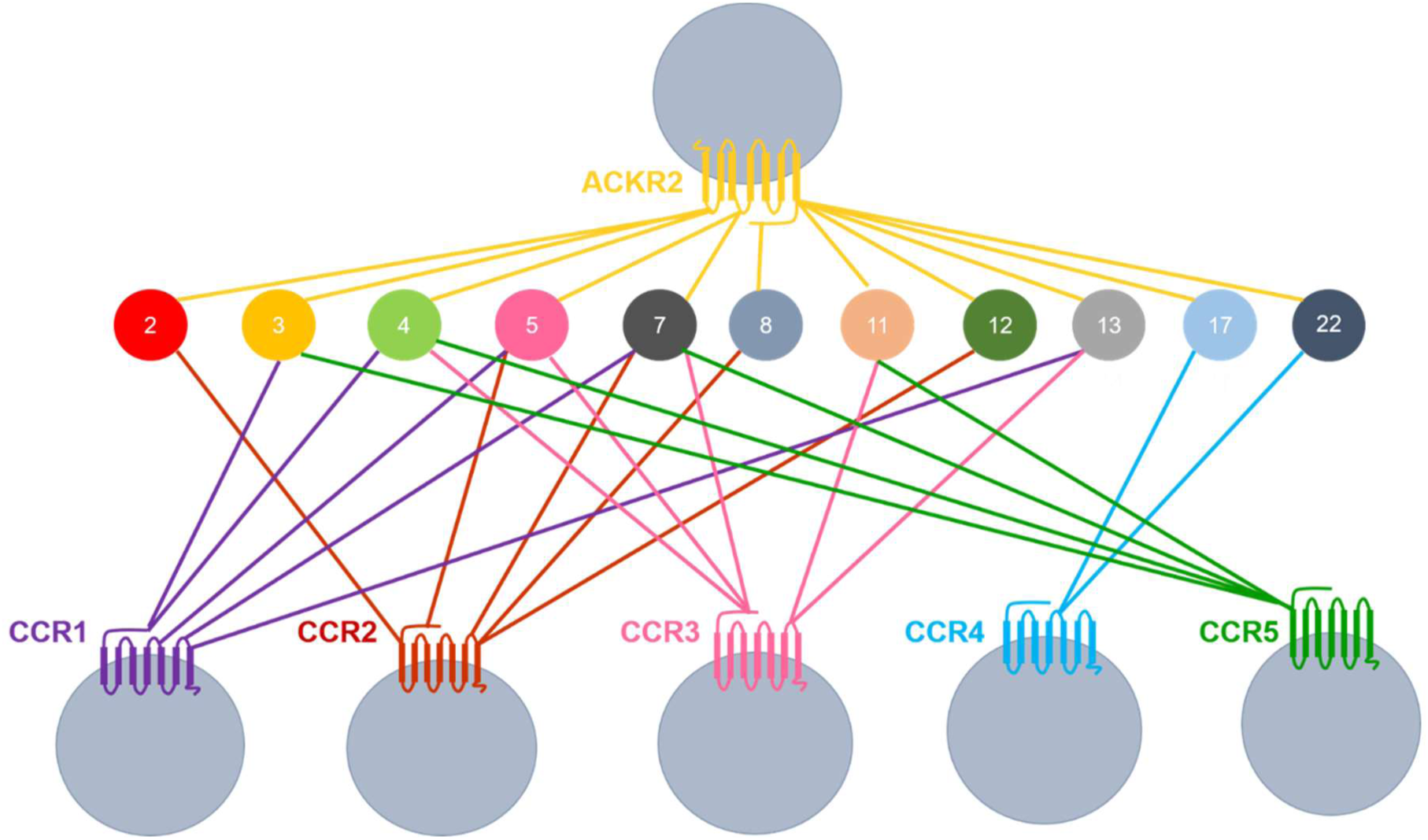
ACKR2 shares ligands with inflammatory chemokine receptors. Coloured lines indicate receptor ligand interactions. Data compiled from Bachelerie, F. *et al.*, 2014 and Nibbs, R. J. B. & Graham, G. J., 2013.

Here we identify CCR1, and its ligand CCL7, as key regulators working with ACKR2 in a reciprocal manner to regulate macrophage numbers, and branching morphogenesis, in the developing mammary gland. Collectively, this study sheds important light on the regulation of macrophage dynamics during virgin mammary gland development.

## Results

### Ductal branching in the pubertal mammary gland is regulated by CCR1

To determine involvement of CCR1, CCR3 and CCR5 in the regulation of ductal branching morphogenesis in the mammary gland we analysed carmine alum stained whole-mounts of mammary glands from 7 week old WT and CCR1-/-, CCR3-/- and CCR5-/- mice (Fig. 2ai-iii). The individual receptor deficient mice have different genetic backgrounds, therefore mice from each strain were compared to their specific WT (Douglas P Dyer *et al.*, 2019). Quantitative analysis of the whole-mounts indicated that branched area, ductal elongation, TEB number and width were unaffected in CCR3-/- and CCR5-/- mice (Fig. 2aii-iii, Supplementary Fig. 1). In contrast, CCR1-/- mice exhibited delayed mammary gland development with decreased branched area at 7 and 8 weeks, reduced ductal elongation and decreased number and width of TEBs at 7 weeks (Fig. 2ai and bi-iv). In addition, in comparison to WT mice, CCR1-/- mice had thinner branches at 8 weeks (Fig. 2bv). This was not seen for CCR3-/- or CCR5-/- mice (Supplementary Fig. 1e). As observed for ACKR2-/- mice, by 12 weeks, when TEBs have regressed and ductal outgrowth is completed, branched area and ductal elongation are equivalent between WT and CCR1-/- mice (Fig. 2bi-ii). Together these data show that CCR1 regulates mammary gland development at a time point coincident with ACKR2 function in the same context.

**Figure 2:**
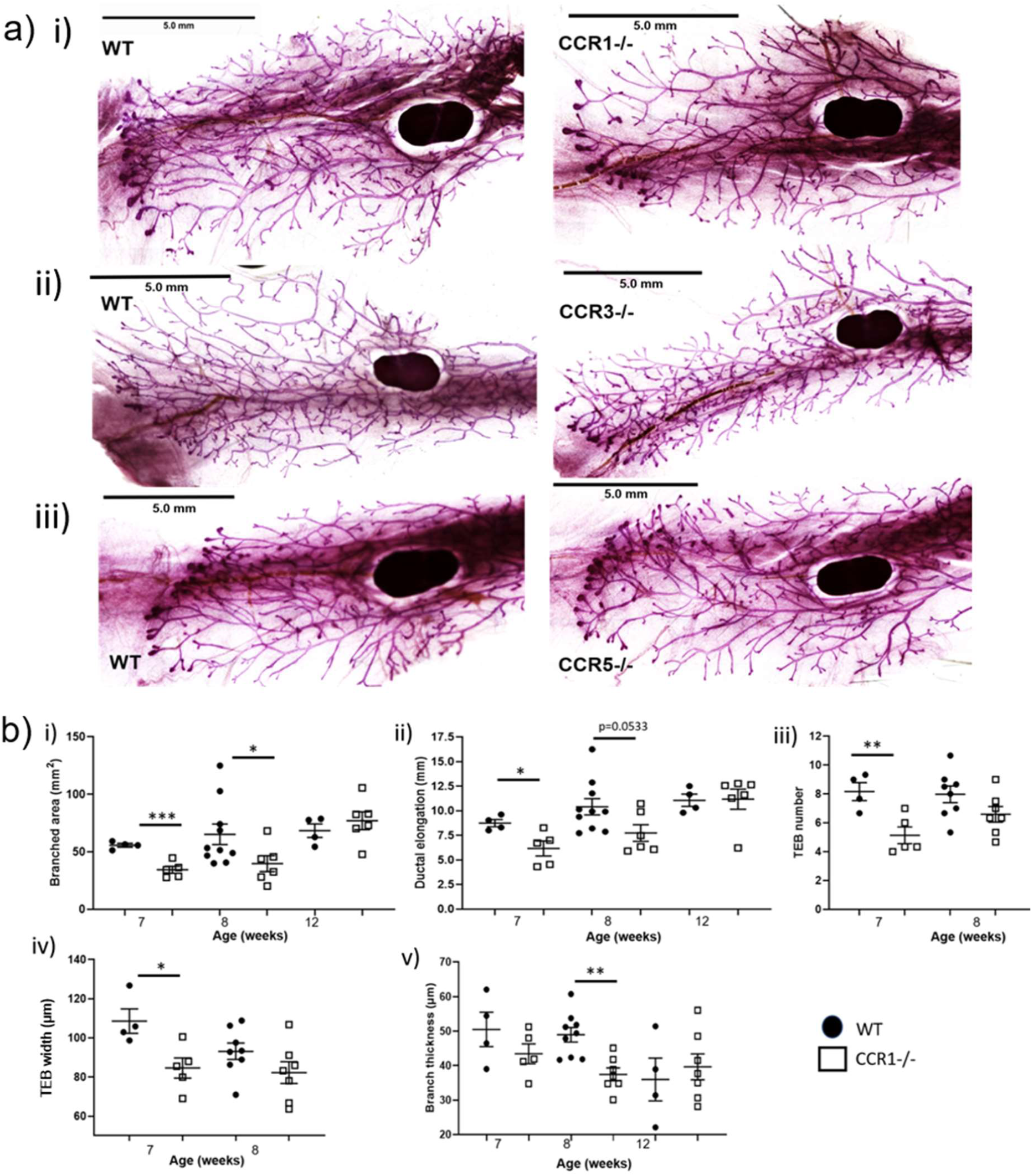
Ductal branching in the pubertal mammary gland is regulated by CCR1. **a)** Representative carmine alum whole mount images of late pubertal (7 week old) virgin mammary glands from **i)** wild-type and CCR1-/-, **ii)** CCR3-/- and **iii)** CCR5-/- mice. Scale bars: 5 mm. **b)** Branching morphogenesis was quantified in 7 (WT n=4, CCR1-/- n= 5), 8 (WT n=10, CCR1-/- n= 7) and 12 (WT n=4, CCR1-/- n= 7) week mammary glands using ImageJ, by measuring: **i)** the area of branching from the inguinal lymph node, and **ii)** ductal elongation, measured from the middle of the inguinal lymph node to the furthest edge of ductal outgrowth. **iii)** The number of TEBs, was determined as the average number from at least 2 individual fields of view (FOV) (5×) per gland. **iv)** The average width of all TEBs was determined from at least 2 F.O.V (5x) per gland. **v)** Branch thickness was determined as the average of 6 measurements from 3 F.O.V (5x) per gland. Significantly different results are indicated. Error bars represent S.E.M.

Of note, in contrast to ACKR2-/- mice, no difference was observed in the distance between, or density of, branches in WT and CCR1-/- mammary glands at any of the time points investigated (Supplementary Fig. 2). This suggests that CCR1 does not regulate the density, but the spread of the ductal network.

Importantly, previous publications have suggested potential redundancy in roles for CCR1, 3 and 5 in vivo (Mantovani, 1999; Schall and Proudfoot, 2011). Whilst we have shown this not to be the case in acute inflammation (Douglas P. Dyer *et al.*, 2019), we have not examined potential receptor redundancy in the context of mammary gland development. Therefore, to test for any potential redundancy between the CCRs, mammary gland whole-mounts were obtained from iCCR-/- mice which have a compound deletion of CCR1, CCR2, CCR3 and CCR5 (Douglas P Dyer *et al.*, 2019). As observed in the absence of CCR1, iCCR-/- mice display similar delayed development at 7 weeks as demonstrated by reduced TEB number (Supplementary Fig. 1c). No additional combinatorial effects of the receptors were observed indicating that CCR1 is a non-redundant regulator of mammary gland development.

### CCR1 and ACKR2 are expressed surrounding epithelium in the mammary gland

We next examined the expression patterns of CCR1 and ACKR2 within the developing mammary gland during late puberty. We used flow cytometry to identify the cell type(s) expressing CCR1 within the mammary gland. As currently available antibodies to murine CCR1 are of limited quality we included cells from CCR1-/- mice as a control. This analysis demonstrated that CCR1 is only detectable on macrophages (CD45+SiglecF- CD11b+F4/80+) within the mammary gland (Fig. 3a) and further in situ hybridisation showed the CCR1+ cells to be intimately associated with the ductal epithelium (Fig. 3b). In contrast to macrophages, eosinophils (CD45+ SiglecF+) and stromal and epithelial (CD45-) cells did not express CCR1 (Fig. 3a). We next examined ACKR2 expression in the mammary gland. Previously, we showed that ACKR2 is expressed by stromal fibroblasts in the developing virgin mammary gland (Wilson *et al.*, 2017). Here we have used in situ hybridisation to locate expression of ACKR2 to stromal cells in the vicinity of the ductal epithelium. Importantly no in situ hybridisation signals were seen in the stroma of CCR1-/- or ACKR2-/- mammary glands (Fig. 3b).

**Figure 3:**
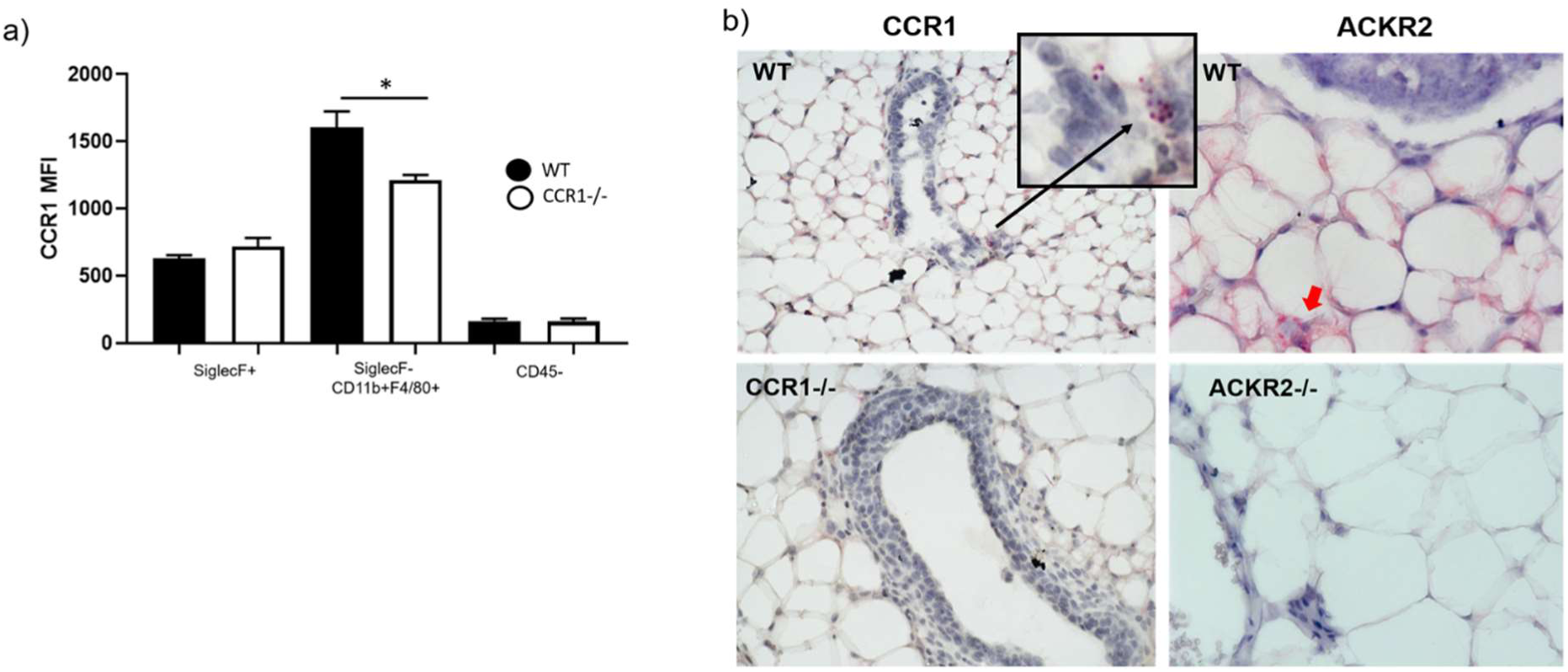
CCR1 and ACKR2 are expressed surrounding epithelium in the mammary gland. **a)** Flow cytometry analysis of CCR1 expression by enzymatically digested WT (black bars, n=6) and CCR1-/- (white bars, n=4) mammary gland cells: CD45-, CD45+ SiglecF+, and CD45+SiglecF-CD11b+F480+. **b)** RNAscope® in situ hybridization of CCR1 (highlighted by black arrow) and ACKR2 (highlighted by red arrow), in the developing virgin mammary gland of WT, CCR1-/- and ACKR2-/- mice. Significantly different results are indicated. Error bars represent S.E.M.

These data therefore demonstrate that CCR1 and ACKR2 are expressed in distinct cell types surrounding TEBs in the developing mammary gland.

### Estrogen induces CCR1 expression on macrophages

We next examined regulation of CCR1 expression on mammary gland macrophages. Estrogen is essential for mammary gland development and ductal epithelial growth and proliferation (Russell C. Hovey, Josephine F. Trott, 2002). ELISA-based analysis of estradiol levels in the plasma of the developing mouse indicated that its production rises over the same time-frame in which we observe altered ductal development in CCR1-/- mammary glands (Fig. 4a). Notably, there was no difference in the levels of estradiol in WT and ACKR2-/- mice, suggesting that the accelerated branching in ACKR2-/- mice is not caused by increased levels of estrogen. To determine whether estrogen regulates CCR1 expression on mammary gland macrophages we enzymatically digested mammary glands and exposed the cells to DMSO (vehicle control) or 17β-estradiol for 1h at 37°C. CCR1 expression was analysed by flow cytometry and shown to increase on CD45+ CD11b+F4/80+ macrophages in response to 17β-estradiol (Fig. 4b). There was no significant difference between the level of CCR1 expression on WT and ACKR2-/- macrophages after exposure, indicating that ACKR2 does not regulate this process.

**Figure 4:**
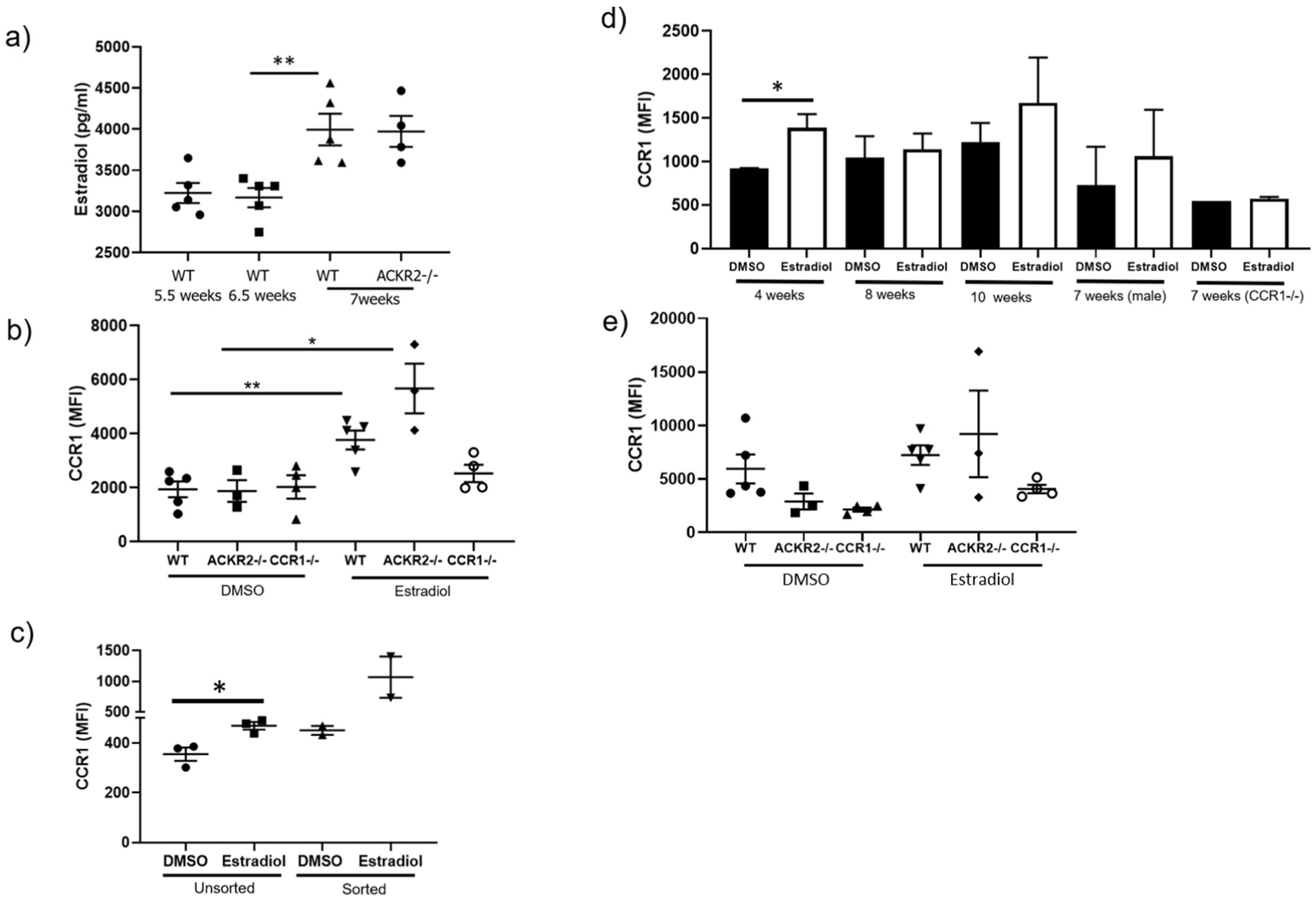
Estrogen induces CCR1 expression on macrophages. **a)** Estradiol levels in plasma from 5.5 weeks WT (n=5), 6.5 weeks WT (n=5), 7 weeks WT (n=5), and 6.5 weeks ACKR2-/- (n=4). **b)** CCR1 expression by CD11b+F4/80+ cells in response to DMSO and 50 μg/ml estradiol in WT (n=5), CCR1-/- (n=4), and ACKR2-/- (n=3). **c)** unsorted and CD11b+F4/80+ FACS sorted cells from the mammary gland. **d)** CCR1 expression by CD11b+F4/80+ cells from female WT mammary glands at 4, 8, 10 weeks old, female 7 week old CCR1-/- mice and male WT inguinal fat pads from 7 week old mice, in response to DMSO and 50 μg/ml 17β-estradiol (each group, n=3); and **e)** CD11b+F4/80+ cells from the peritoneum. Significantly different results are indicated. Error bars represent S.E.M.

To determine whether this was a direct effect of estradiol on mammary gland macrophages, CD11b+F4/80+ cells were isolated by FACS. In the absence of other cell types, CCR1 expression was increased following exposure to 17β-estradiol indicating that estrogen induction of CCR1 results from a direct effect on mammary gland macrophages (Fig. 4c).

Notably, upregulation of CCR1 on macrophages in response to estradiol is age dependent, as there is no difference in CCR1 expression in mice older than 8 weeks (Fig. 4d). In addition, 17β-estradiol has no effect on macrophages isolated from the male fat pad or the peritoneum of pubertal female mice (Fig. 4d-e). Taken together, this suggests that the effect of estrogen on CCR1 expression is restricted to pubertal mammary gland macrophages and limited to the key developmental time-frame we have identified.

### Chemokine levels are altered in the absence of CCR1 and ACKR2

To identify the specific chemokines involved in regulating mammary gland development through CCR1 and ACKR2, multiplex protein analysis of mammary gland lysates was carried out. In keeping with our previous data we showed that, in the absence of scavenging by ACKR2, the chemokines CCL7, CCL11 and CCL12 accumulate in the mammary gland at 6-7 weeks (Fig. 5a) (Wilson *et al.*, 2017). The current analysis further revealed elevated CCL3, CCL19, CCL22 and CXCL10 in the ACKR2 -/- mammary gland over this time-frame (Fig. 5a). Notably, other key chemokines associated with monocyte and macrophage migration i.e. CCL2 and CCL5 are unchanged in the ACKR2-/- mammary gland (Fig. 5a). Importantly, there were no significant differences in the levels of these chemokines in lysates obtained from male WT and ACKR2-/- inguinal fat pads, indicating that the changes observed in female lysates are specifically associated with the mammary gland (Supplementary Fig. 3). In CCR1-/- mice, the levels of CCL7, CCL11 and CCL12 were unchanged, indicating that ACKR2 is functional in these mice and able to scavenge chemokines normally. CCL3, CCL19, CCL22, CXCL1 and CXCL12 are increased in ACKR2-/- mice and decreased in CCR1-/- mice (Fig. 5). Given that CCL19, CXCL1 and CXCL12 are not ligands for either ACKR2 or CCR1, it is likely that, along with CCL3 and CCL22, their altered levels reflect variation in the numbers of chemokine-expressing immune or epithelial cells within the mammary gland.

**Figure 5:**
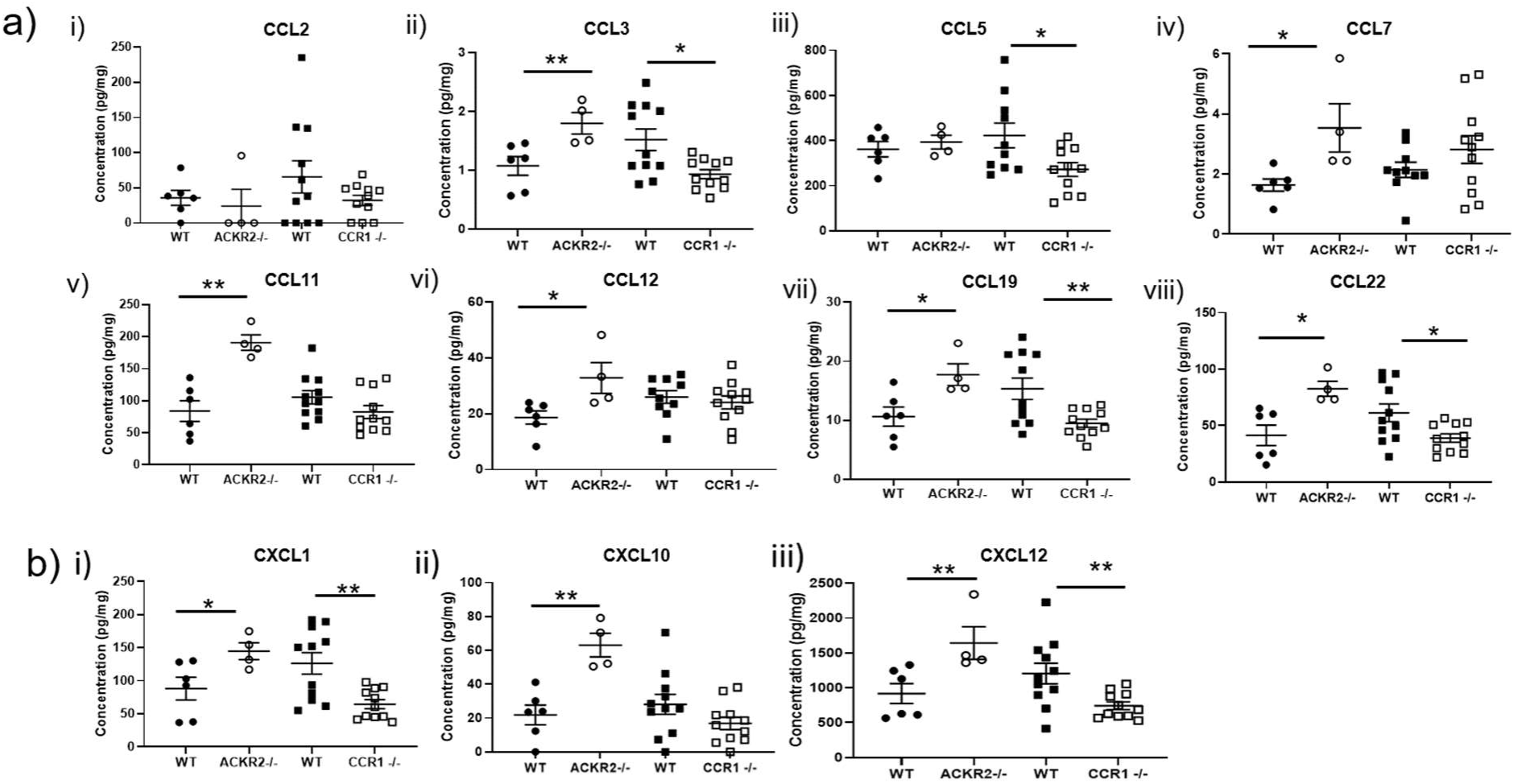
Chemokine levels are altered in the absence of ACKR2 and CCR1. Multiplex measurement of protein concentration of **a)** inflammatory CC-chemokines; **i)** CCL2, **ii)** CCL3, **iii)** CCL5, **iv)** CCL7, **v)** CCL11, **vi)** CCL12, **vii)** CCL19 and **viii)** CCL22, and; **b)** inflammatory CXC chemokines **i)** CXCL1, **ii)** CXCL10 and **iii)** CXCL12 in whole mammary gland homogenates. WT (CCR1) n=11, CCR1-/- n=11, WT (ACKR2) n=6, and ACKR2-/- n=4. Significantly different results are indicated. Error bars represent S.E.M.

### CCR1 and ACKR2 reciprocally regulate CD206+ macrophages within the mammary gland

Reciprocal regulation of leukocyte dynamics by CCR1 and ACKR2 in the developing mammary gland should be reflected in complimentary changes in levels of key cellular populations in CCR1-/- and ACKR2-/- mice. We detected no significant differences in the lymphocyte populations or in non-macrophage myeloid cell populations investigated. However, differences in a key macrophage population were identified. To investigate the effects of CCR1 deficiency on macrophage levels in the mammary gland, flow cytometry of enzymatically digested 6.5 week old WT and CCR1-/- glands was carried out. The gating strategy employed is described in Supplementary Fig. 4. CCR1-/- mice displayed no significant differences in the bulk macrophage population (CD45+CD11b+F4/80+) (Fig. 6ai and bi). However, we detected a significant decrease in the percentage of a small population of macrophages expressing CD206 (mannose receptor) (CD45+SiglecF-F4/80+CD206+) in CCR1-/- mice (Fig. 6aii, bii). Analysis of ACKR2-/- mice revealed a complimentary phenotype to CCR1-/- mice in that they displayed an increase in the percentage of macrophages in the mammary gland population and specifically of the CD206+ macrophage subset (Fig. 6cii).

**Figure 6:**
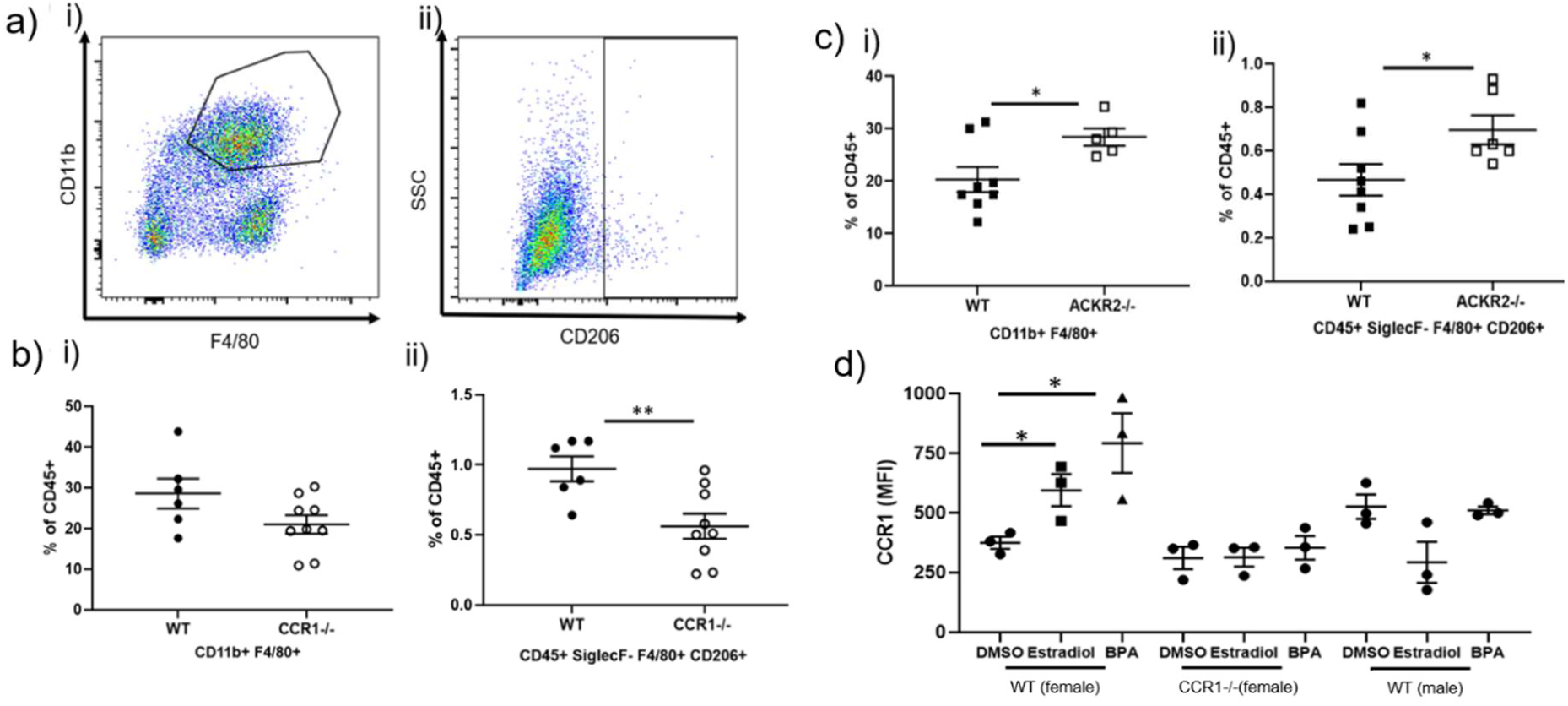
CCR1 and ACKR2 reciprocally regulate CD206+ macrophages within the mammary gland. **a)** Flow cytometry was used to determine the percentage of i) CD11b+F4/80+ and ii) SiglecF- F4/80+ CD206+ macrophages, within the CD45+ compartment of the 6.5 weeks old developing mammary gland. Flow cytometry of **b)** WT (n=6) and CCR1-/- (n=9) and **c)** WT (n=8) and ACKR2-/- (n=5) mammary gland cells was carried out. **d)** CCR1 expression by SiglecF- F4/80+ CD206+ cells in response to DMSO and 50 μg/ml 17β-estradiol and Bisphenol A; female WT and CCR1-/- and male WT (n=3). Significantly different results are indicated. Error bars represent S.E.M

Finally, we examined the effects of estrogen on the CD206+ macrophage population. Our data show that CCR1 expression was also increased on the surface of CD206+ macrophages in response to both 17β-estradiol and the estrogen mimic Bisphenol A (BPA) (Fig. 6d). No effect of estrogen on CCR1 expression was observed in male macrophages (Fig. 6d).

Thus, a key population of CD206+ macrophages are reciprocally regulated by ACKR2 and CCR1. Importantly, CD206+ mammary gland macrophages have previously been implicated in branching morphogenesis (Jäppinen *et al.*, 2019) and we propose that ACKR2 and CCR1 reciprocally control this population to coordinate branching morphogenesis in the pubertal mammary gland.

### CCL7 regulates CD206+ macrophages and branching morphogenesis

Of the chemokines detected within the mammary gland, CCL7 is of particular interest as it is shared between CCR1 and ACKR2 (Fig. 1), and is elevated in the pubertal mammary glands of ACKR2-/- mice (Fig. 5aiv)(Wilson *et al.*, 2017). In addition, qRT-PCR analysis also revealed that CCL7 is transcribed, by purified F4/80+ cells, at higher levels than other ACKR2 ligands (Fig. 7ai). We therefore investigated its expression and function in the mammary gland. Using flow cytometry, intracellular staining revealed that CCL7 is produced by immune cells, including SiglecF+ eosinophils, SiglecF- F4/80+ macrophages, and SiglecF-Ly6C+ monocytes (Fig. 7aii). For each cell type, a markedly higher percentage of cells obtained from the female mammary gland produced CCL7, than from male fat pad cells. Notably, around 60% of female SiglecF+ cells produced CCL7 compared with 10% of male cells (Fig. 7aii). The percentage of CCL7+ cells was unaffected in the absence of ACKR2 (Fig. 7ai, ii). CCL7 is also produced by CD45- epithelial cells: mature (EpCAM+ CD49f-) and progenitor luminal (EpCAM+ CD49f+), and basal (EpCAM - CD49f+) cells (Fig. 7aiii, Supplementary Fig. 4b). Further, bioinformatic analysis confirmed that CCL7 is produced by epithelial cells, including basal, luminal and myoepithelial cells (Supplementary Fig. 5) (Bach *et al.*, 2017).

**Figure 7:**
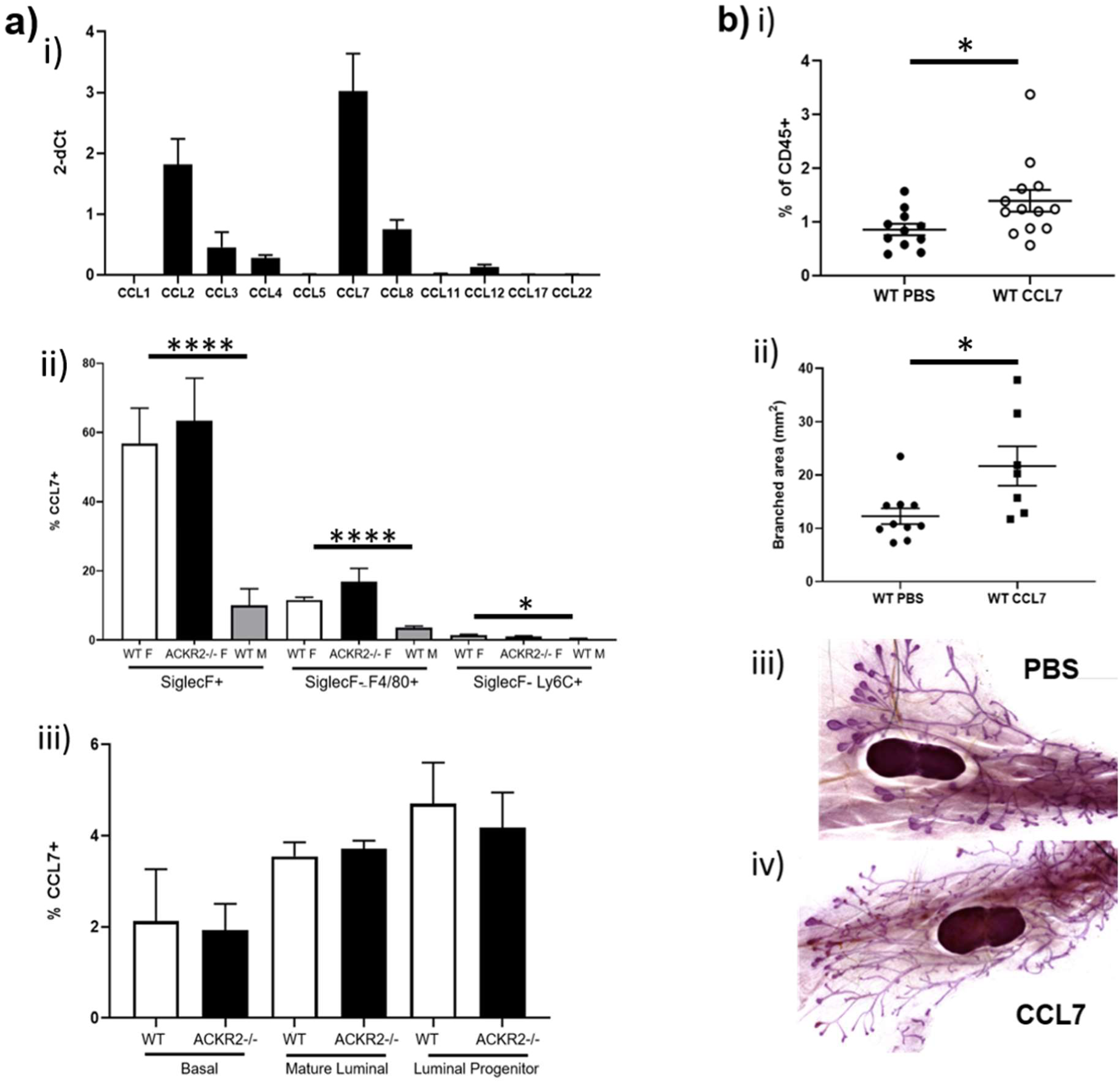
CCL7 controls CD206+ macrophages and branching morphogenesis. **a)** CCL7 is produced in the mammary gland by **i)** immune cells; including SiglecF+, SiglecF- F4/80+ and SiglecF-Ly6C+, male WT (n=7) and female WT (n=6) and ACKR2-/- (n=4). **ii)** Transcription of inflammatory chemokines by purified F4/80+ cells (n=3) and **iii)** CCL7 production by epithelial cells, Mature Luminal (EpCAM+ CD49f-), progenitor luminal (EpCAM+ CD49f+), and basal (EpCAM- CD49f+), female WT (n=6) and ACKR2-/- (n=4). **b)** 3 days after subcutaneous administration of PBS or 2 μg CCL7 at 6 weeks, **i)** the percentage of SiglecF-F4/80+CD206+ cells was measured by flow cytometry. (PBS, n=11, CCL7, n=13) and **ii)** the area of branching was measured using Image J (PBS, n=10, CCL7, n=7). **iii)** and **iv)** Representative images of whole mounts from PBS and CCL7 injected mice. Significantly different results are indicated. Error bars represent S.E.M.

Given the notable CCL7 expression in the mammary gland, we next directly tested its potential role in mammary gland development. PBS or 2 μg of CCL7 was administered subcutaneously at the site of the mammary fat pad at the key time point of 6 weeks. After 3 days, mammary glands were harvested for cellular analysis by flow cytometry and carmine alum whole-mount analysis. CCL7 administration alone was sufficient to increase the percentage of CD206+ macrophages, and the area of branching within the mammary gland (Fig. 7b). These data confirm that elevated levels of CCL7, as observed in ACKR2-/- mice, leads to increased numbers of CD206+ macrophages in the mammary gland and accelerated branching.

Overall these data demonstrate a role for CCL7, a ligand shared by CCR1 and ACKR2, in branching morphogenesis. Lending further support to this conclusion is the fact that bioinformatic interrogation of the precocious puberty (CTD Gene-Disease Associations) dataset, using Harmonizome (Rouillard *et al.*, 2016), revealed that CCL7 and ACKR2 are both associated with precocious puberty in children, with standardized values of 1.25588 (p=0.09) and 1.02634 (p=0.011) respectively.

## Discussion

The importance of macrophages in controlling developmental processes is well known (Wynn, Chawla and Pollard, 2013). The role of chemokines and their receptors, which provide molecular cues to guide and position macrophages during development, is an emerging area of research (Lee *et al.*, 2014; Wilson *et al.*, 2017). Previously, we revealed that the scavenging atypical chemokine receptor, ACKR2 controlled macrophages in the mammary gland through a CCR2-independent pathway (Wilson *et al.*, 2017). Here we have revealed a previously unknown immunological mechanism whereby, ACKR2 and the inflammatory chemokine receptor CCR1, interact with their shared ligand CCL7 to coordinate the levels of CD206+ macrophages, and thus, the extent of branching morphogenesis in the pubertal mammary gland. Importantly, administration of CCL7 alone was able to increase the percentage of CD206+ macrophages within the mammary gland and drive accelerated branching morphogenesis. We propose that in CCR1-/- mice, although CCL7 levels are unaltered, macrophages are unable to sense and respond to the ligand without the cognate receptor, leading to delayed branching (Fig. 8).

**Figure 8:**
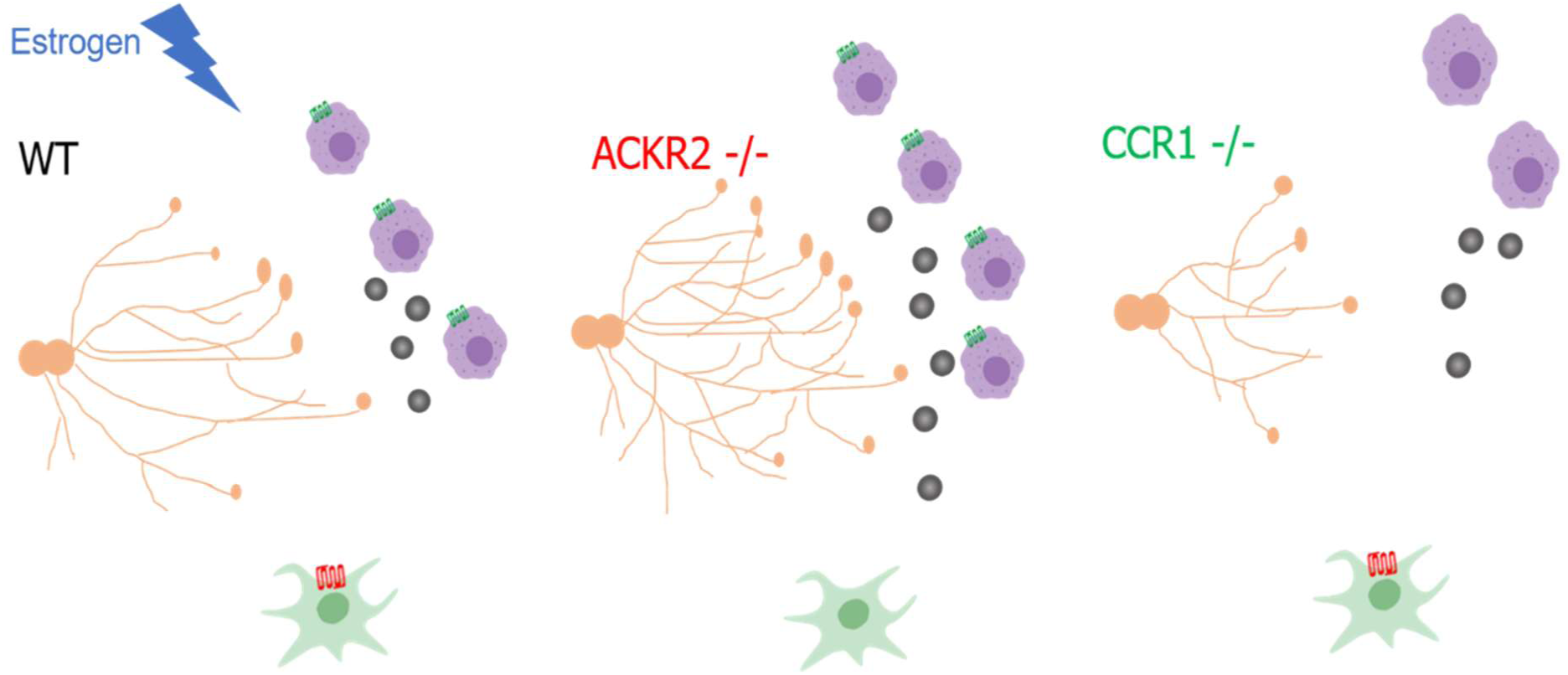
Proposed mechanism by which chemokine receptors CCR1 and ACKR2 coordinate mammary gland development. Estrogen increases CCR1 expression on macrophages (purple) during puberty, and stromal fibroblast (green) expressed ACKR2 modulates levels of CCL7 (grey circles) to control the movement of CCR1+ macrophages to the ductal epithelium (orange). Schematic image was created with BioRender.

Previously, it was thought that all mammary gland macrophages, at rest, and in pathology, were derived from the bone marrow (Coussens and Pollard, 2011). In our previous study, we showed that branching was unaltered in the absence of CCR2, indicating that the macrophage population responsible for promoting branching morphogenesis was unlikely to be bone marrow derived (Wilson *et al.*, 2017). Recently, a novel CD206+ macrophage population has been identified in the mammary gland, which is unaffected in the absence of CCR2, but reduced in *plvap*-/- mice, which have reduced numbers of foetal-derived macrophages (Jäppinen *et al.*, 2019). Branching is severely impaired in these mice suggesting that foetal-derived macrophages play a key role in promoting branching morphogenesis (Jäppinen *et al.*, 2019). We believe that the macrophage population identified in our study may be derived from the same embryonic population (Jäppinen *et al.*, 2019).

CCR1 is an inflammatory chemokine receptor which is expressed by immune cells, and has been shown to be important in a number of pathologies, including: sepsis, viral infections, cancer and autoimmune disease (Domachowske *et al.*, 2000; Katschke Jr. *et al.*, 2001; Ness *et al.*, 2004; Kitamura *et al.*, 2015). To our knowledge, this is the first description of a key role for CCR1 in development. Of note, in the placenta, CCR1 has been shown to be expressed by human trophoblasts as they switch to an invasive phenotype (Sato *et al.*, 2003). ACKR2 is highly expressed by placental trophoblasts, preventing excess levels of inflammatory chemokines from entering the foetus, from the mother’s circulation, by a process of chemokine compartmentalisation (Teoh *et al.*, 2014; Lee *et al.*, 2019). As CCR1 expression has also been described in placental development, there could be wider implications of the interaction described in this study.

In the mouse, sexual maturity occurs at around 6 weeks (Topper and Freeman, 1980). Here we report a marked increase in plasma estradiol levels between 6.5 and 7 weeks. This is the key time point in ACKR2/CCR1-dependent regulation of branching morphogenesis. ACKR2 expression in the mammary gland specifically peaks at 6.5 weeks and branching begins to accelerate at this time point (Wilson *et al.*, 2017). We show that 17β-estradiol increases CCR1 expression on macrophages. However, this is restricted to pubertal mammary gland macrophages, as older female, male and peritoneal macrophages do not respond. In addition to 17β-estradiol, the estrogen mimic, Bisphenol A also increased CCR1 expression on CD206+ macrophages. This may be of concern as BPAs are widely found in the environment and could potentially alter the immune response, and the extent of branching in the mammary gland, in children during puberty. Previously CCR1 expression on T cells was shown to be regulated by 17β-estradiol (Mo *et al.*, 2005). However this is the first description of estrogen controlled CCR1 expression on macrophages. This observation could have implications for our understanding of diseases where females exhibit increased susceptibility. One example is rheumatoid arthritis, where CCR1 is also associated with pathology (Katschke Jr. *et al.*, 2001; van Vollenhoven, 2009).

Understanding the molecular signals which guide the rate of branching morphogenesis in the mammary gland is highly important. Precocious puberty is a condition where puberty begins before the age of 8, with some girls developing breasts as early as 4. This results from early activation of the gonadotropic axis, leading to accelerated growth and bone maturation, but ultimately reduced stature (Carel *et al.*, 2004). Potential risk factors include exposure to endocrine disrupters, obesity, stress and ethnicity (Cesario and Hughes, 2007; Lee *et al.*, 2007; Meeker, 2012; Kelly *et al.*, 2017). As mammary gland development is delayed in mice in the absence of CCR1, this could represent a novel therapeutic target to treat aspects of precocious puberty. Several CCR1 antagonists are available and have been used in a number of clinical trials (Lebre *et al.*, 2011). In addition, early breast development leads to higher risks of breast cancer in later life (Bodicoat *et al.*, 2014), and women with dense breasts are more likely to develop breast cancer (Nazari and Mukherjee, 2018). This can be related to poor detection by mammography as the branches mask the cancer, but may also be caused by genetic factors, parity and alterations in the breast stroma. Both ACKR2 and CCR1 have been shown to be important in the progression of breast cancer, therefore understanding early interactions between these receptors could reveal key insights, which drive later pathology (Kitamura *et al.*, 2015; Shin *et al.*, 2017; Hansell *et al.*, 2018).

In this study, we have uncovered a novel mechanism by which estradiol upregulates CCR1 expression by pubertal mammary gland macrophages and stromal ACKR2 modulates levels of CCL7, to control the movement of the CCR1+ macrophages to the ductal epithelium. Overall therefore our data demonstrate that CCR1 and ACKR2 coordinately regulate mammary gland branching morphogenesis.

## Methods

### Animals

Animal experiments were carried out under the auspices of a UK Home Office Project Licence and conformed to the animal care and welfare protocols approved by the University of Glasgow. C57BL/6 mice, ACKR2-/- (Jamieson *et al.*, 2005), CCR1-/-, CCR3-/-, CCR5-/- and iCCR-/- (Douglas P. Dyer *et al.*, 2019) mice were bred at the specific pathogen-free facility of the Beatson Institute for Cancer Research.

### Carmine Alum Whole Mount

Carmine alum whole mounts were carried out as described previously (Wilson *et al.*, 2017). Briefly, fourth inguinal mammary glands were fixed overnight in 10% neutral buffered formalin (NBF) (Leica) at 4°C. Glands were dehydrated for 1 h in distilled water, followed by 70% ethanol and 100% ethanol before overnight incubation in xylene (VWR international). Tissue was rehydrated by 1 h incubation in 100% ethanol, 70% ethanol and distilled water, before staining in Carmine Alum solution overnight at room temperature (0.2% (w/v) carmine and 10 mM aluminium potassium sulphate (Sigma)). Tissue was dehydrated again before overnight incubation in xylene. Finally, glands were mounted with DPX (Leica) and stitched bright-field images at 10× magnification were taken using an EVOS FL auto2 microscope (Thermofisher). Ductal elongation, and branched area from the lymph node, were measured using ImageJ 1.52a (Schneider, Rasband and Eliceiri, 2012). 5 x brightfield images were obtained using the Zeiss Axioimager M2 with Zen 2012 software. The numbers of branches and branch thickness were counted as the average from 3 measurements from 6 individual fields of view (F.O.V.) from each whole mount. TEBs were counted as the average from at least 2 F.O.V. from each whole mount. All samples were blinded before measurements were taken.

### RNAscope ® In situ hybridisation

Mammary glands were fixed in 10% neutral buffered formalin at room temperature for 24-36 hours before being dehydrated using rising concentrations of ethanol and xylene, and paraffin embedded (Shandon citadel 1000 (Thermo Shandon). Tissue was sectioned onto Superfrost plus slides (VWR) at 6 μm using a Microtome (Shandon Finesse 325 Microtome, Thermo). Slides were baked at 60°C for 1 h before pre-treatment. Slides were deparaffinised with xylene (5 mins x 2) and dehydrated with ethanol (1 min x 2). Tissues were incubated with Hydrogen peroxide for 10 mins at RT, then boiled in antigen retrieval buffer for 15 mins. Slides were treated with protease plus for 30 mins at 40°C. Slides were then hybridised using the RNAScope® 2.5 Red Manual Assay (Advanced cell diagnostics) according to the manufacturer’s instructions using the Mm-Ccr1 and Mm-ACKR2 probes. Slides were mounted in DPX (Sigma Aldrich) and imaged on an EVOS FL Auto2microscope.

### Mammary gland digestion

The inguinal lymph node was removed from the fourth inguinal mammary gland, tissue was chopped, and enzymatic digestion was carried out in a 37°C shaking incubator at 200 rpm for 1 h, with 3 mg/ml collagenase type 1 (Sigma) and 1.5 mg/ml trypsin (Sigma) in 2 ml Leibovitz L-15 medium (Sigma). The suspension was shaken for 10 s before addition of 5 ml of L-15 medium supplemented with 10% foetal calf serum (Invitrogen) and centrifugation at 400 g for 5 min. Red blood cells were lysed using Red Blood Cell Lysing Buffer Hybri-Max (Sigma) for 1 min and washed in PBS. Cells were washed in PBS with 5 mM EDTA, resuspended in 2 ml 0.25% Trypsin-EDTA (Sigma) and incubated at 37°C for 2 min before addition of 5 ml of serum-free L-15 containing 1 μg/ml DNase1 (Sigma) for 5 min at 37°C. L-15 containing 10% FCS was added to stop the reaction and cells were filtered through a 40 μm cell strainer before a final wash in FACS buffer (PBS containing 1% FCS and 5 mM EDTA).

### Flow cytometry

Antibodies were obtained from BioLegend and used at a dilution of 1:200: CD45 (30-F11), CD11b (M1/70), F4/80 (BM8), SiglecF (S17007L), Ly6C (HK1.4), EpCAM (G8.8), CD49f(GoH3), CCR1 (S10450E), and CD206 (C068C2) for 30 min at 4°C. Dead cells were excluded using Fixable Viability Dye eFluor 506 (Thermo Fisher). Intracellular staining for CCL7 was carried out using 1 in 100 biotinylated CCL7 antibody (R&D Systems) and Strepdavidin BV605 (BioLegend) and eBioscience intracellular fixation and permeabilization buffer. Flow cytometry was performed using an LSRII or Fortessa, (BDBiosciences) and analysed using FlowJo V10.

### Proteomic analysis

The inguinal lymph node was removed from the fourth inguinal mammary gland, tissue was chopped, frozen in liquid nitrogen, crushed with a mortar and pestle, and resuspended in dH2O containing protease inhibitors (Pierce). Protein levels were determined using a custom designed Magnetic Luminex Multiplex assay (R&D Systems), as described in the manufacturer’s instructions, and read with a Bio-Rad Luminex-100 machine. Data was normalised to the protein concentration of tissue samples, determined by a BCA assay (Pierce).

### Subcutaneous administration of CCL7

2 μg of CCL7 in 200 μl PBS (R&D Systems) was injected subcutaneously into mice at 6 weeks of age. After 3 days, mice were culled and mammary glands were excised and processed for whole mount and cellular analysis.

### 17β-estradiol assays

Fourth inguinal mammary glands were digested to obtain single cell suspensions. Cells were plated at 0.5-1 × 10^5^ cells in a 96 well plate in L-15 media containing 5% FCS and exposed to DMSO (vehicle control) or 50 ug/ml 17-β estradiol or Bis-phenol A (Sigma) for 1 h at 37°C, 5% CO2. The level of 17-β estradiol in plasma samples was determined using the Estradiol parameter kit (R&D Systems) as described in the manufacturer’s instructions.

### Bioinformatic analysis

Chemokine expression by epithelial cells was determined by searching the data repository from Bach et al, 2017 (Bach *et al.*, 2017) at: https://marionilab.cruk.cam.ac.uk/mammaryGland/.

### Statistical analysis

Data were analysed using GraphPad Prism 8.1.2. Normality was assessed using Shapiro Wilk and Kolmogorov–Smirnov tests. For data with normal distribution, two-tailed, unpaired t-tests were used. Where data was not normally distributed, Mann–Whitney tests were used. Significance was defined as p<0.05 *. Error bars indicate standard error of the mean (S.E.M.).

## Acknowledgements

We thank the University of Glasgow’s animal facility staff for the care of our animals and flow cytometry facility staff for technical assistance. The study was supported by a Programme Grant from the Medical Research Council (MR/M019764/1). Work in GJG’s laboratory is also funded by a Wellcome Trust Investigator Award (099251/Z/12/Z). GJG is a recipient of a Wolfson Royal Society Merit award.

## Author Contributions

GJW conceived the study, performed experiments, analysed data and wrote the paper. AF performed experiments and analysed data. SRL, JK, MP and AJH performed experiments. GJG conceived the study and wrote the paper.

