## Supplemental Information for "Chemokine receptors ACKR2 and CCR1 coordinate macrophage dynamics and mammary gland development"

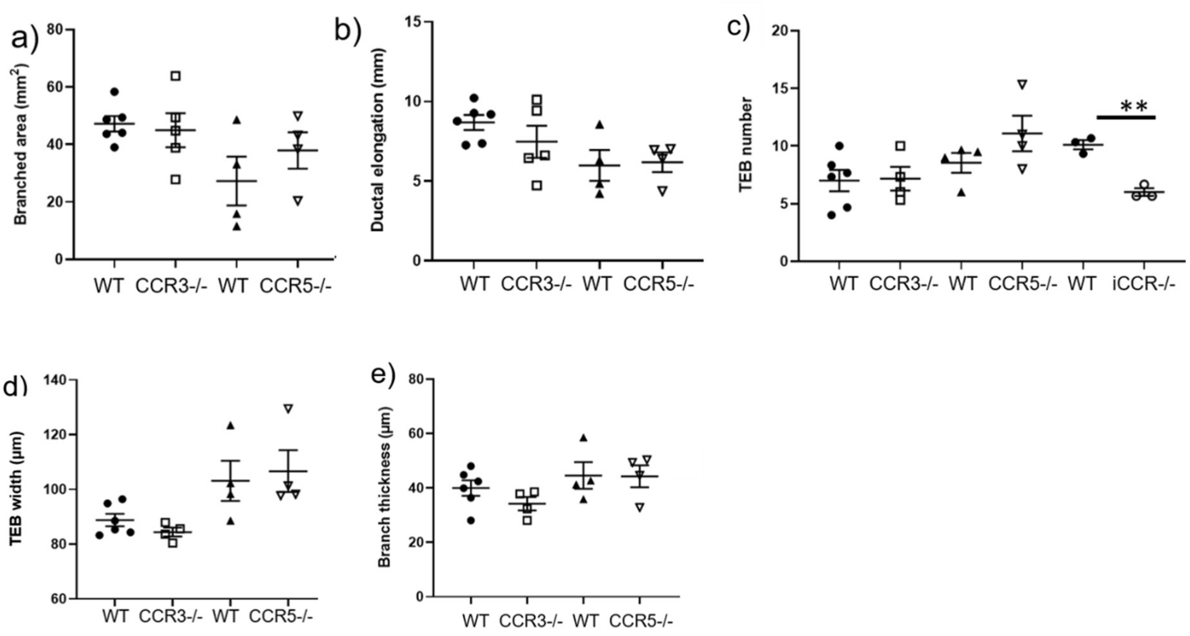


**Supplementary Figure 1: CCR3 and CCR5 do not control branching morphogenesis.** Branching morphogenesis in carmine alum whole mounts were quantified in 7 week mammary glands; (WT (CCR3) n=6, CCR3-/- n= 5; WT (CCR5) n=4, CCR5-/- n= 4). **a)** the area of branching from the inguinal lymph node, **b)** ductal elongation, measured from the middle of the inguinal lymph node to the furthest edge of ductal outgrowth. **c)** The number of TEBs, was determined as the average number from at least 2 individual fields of view (FOV) (5×) per gland WT (iCCR) n=3, iCCR-/- n= 3). **d)** The average width of all TEBs was determined from at least 2 F.O.V (5x) per gland. **e)** Branch thickness was determined as the average of 3 measurements from 6 x F.O.V (5x) per gland. Error bars represent S.E.M.

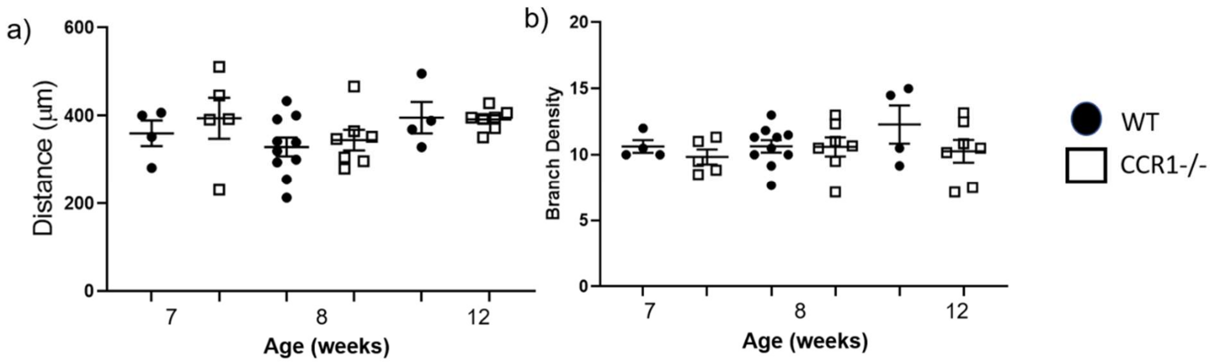


**Supplementary Figure 2: Branch density is unaffected in CCR1-/- mice**. Branching morphogenesis in carmine alum whole mounts were quantified in 7 (WT n=4, CCR1-/- n= 5), 8 (WT n=10, CCR1-/- n= 7)and 12 (WT n=4, CCR1-/- n= 7) week mammary glands; in terms of **a)** the distance between branches, and **b)** the number of branches in a 5 x F.O.V. Each data point represents the average of 3 measurements from 6 individual F.O.V. per gland. Error bars represent S.E.M.

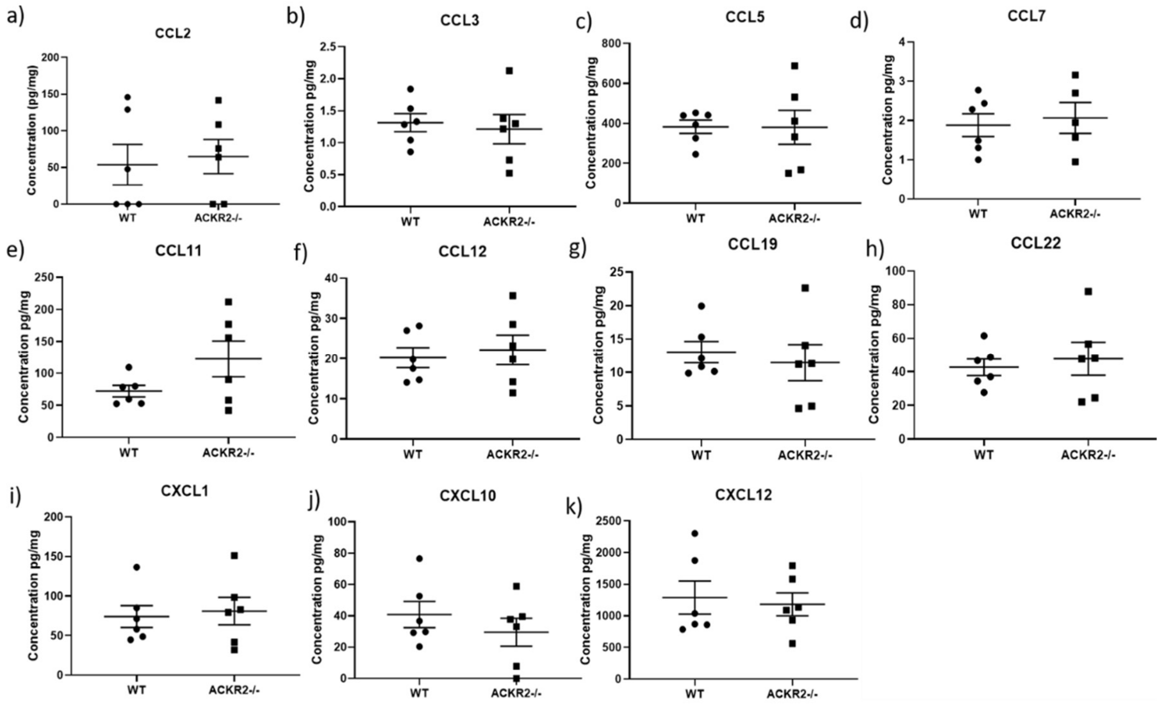


**Supplementary Figure 3: Chemokine levels in the male fat pad are unaffected in the absence of ACKR2.** Multiplex measurement of protein concentration of **a)** CCL2, **b)** CCL3, **c)** CCL5, **d)** CCL7, **e)** CCL11, **f)** CCL12, **g)** CCL19, **h)** CCL22, **i)** CXCL1, **j)** CXCL10 and **k)** CXCL12 in whole fat fad homogenates. WT n=6, and ACKR2-/- n=6. Error bars represent S.E.M.

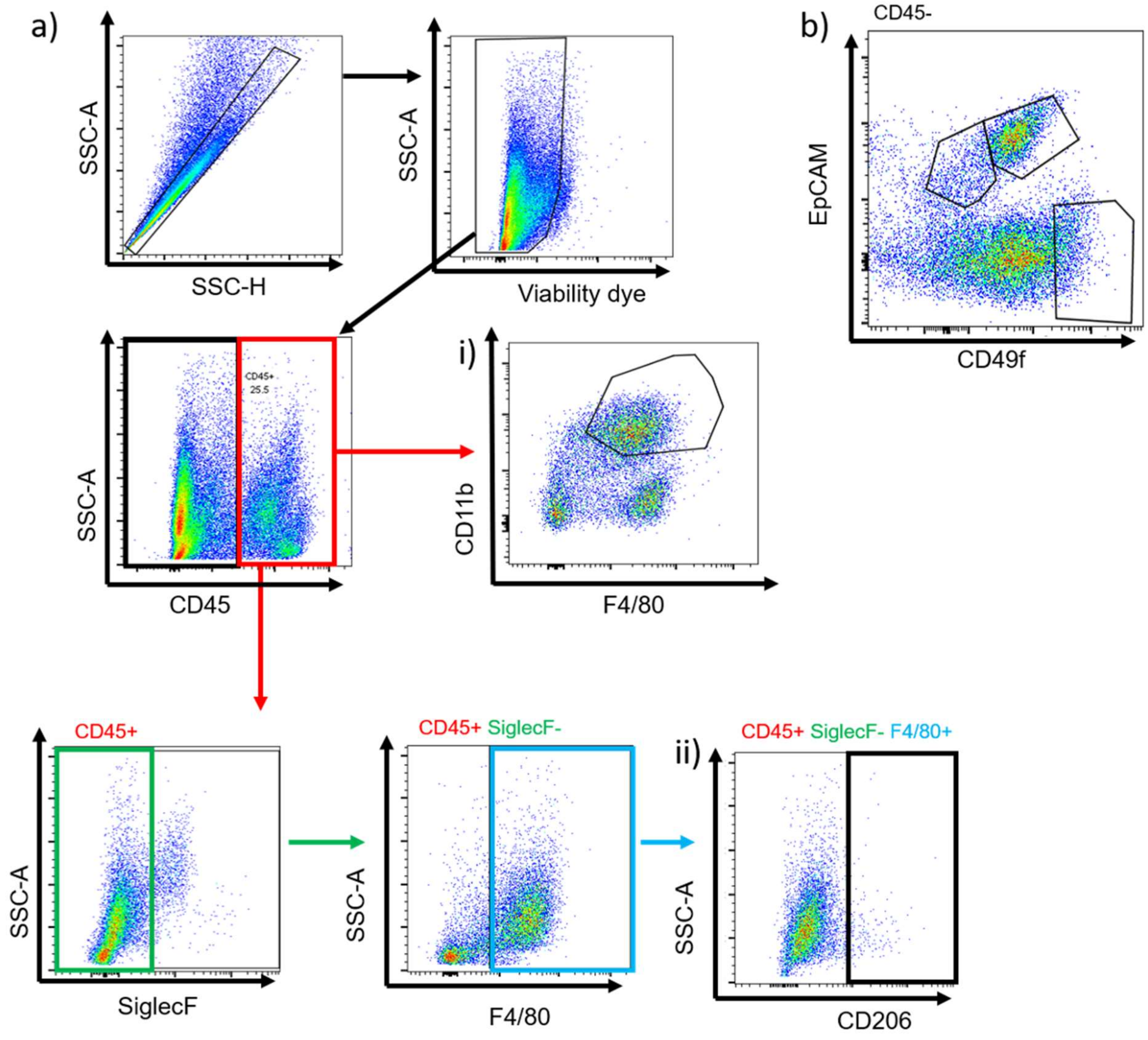


**Supplementary Figure 4: Gating strategy to define immune and epithelial cells in the mammary gland.** Flow cytometry was carried out to measure the percentage of cells in the mammary gland. Initially, single cells were gated, dead cells were excluded, and CD45+ immune cells were gated. Populations were then expressed as a percentage of CD45+ cells, including; **a)** i) CD11b+F4/80+, and ii) SiglecF-F4/80+CD206+ cells. **b)** For epithelial cell subsets live, single CD45- cells were gated. Populations were then expressed as a percentage of CD45- cells, including mature (EpCAM+ CD49f-) and progenitor luminal (EpCAM+ CD49f+), and basal (EpCAM - CD49f+) cells.

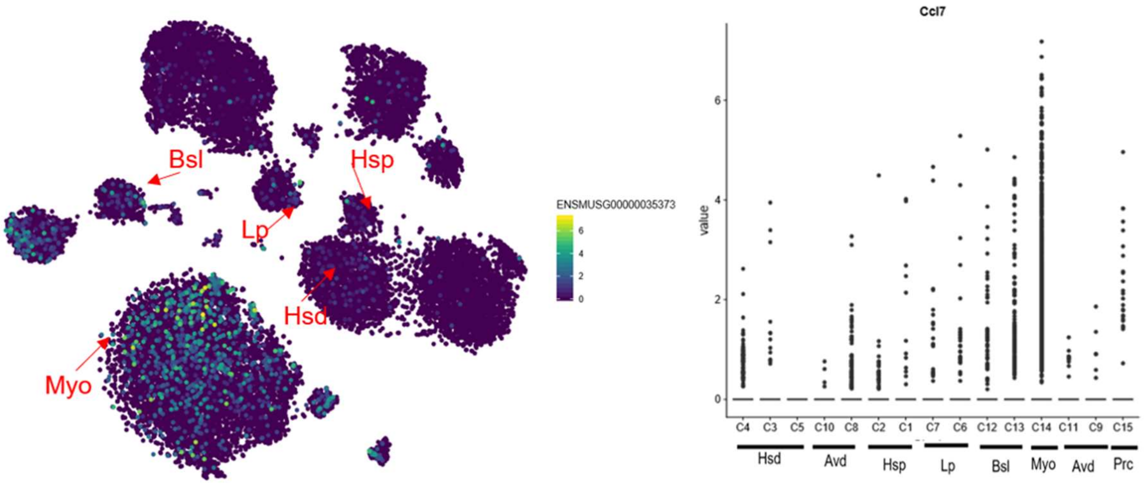


**Supplementary Figure 5: CCL7 is produced by epithelial cell subsets.** Expression of CCL7 by epithelial cells was determined by searching the data repository from Bach *et al*, 2017 [23]at: https://marionilab.cruk.cam.ac.uk/mammaryGland/. Epithelial subsets include; hormone sensing differentiated (Hsd), differentiated alveolar (Avd), hormone sensing progenitor (Hsp), luminal progenitor (Lp), basal (Bsl), myoepithelium (Myo), Procr+ (Prc).
